# Evolution of pineal non-visual opsins in lizards and the tuatara (Lepidosauria)

**DOI:** 10.1101/2024.11.13.623426

**Authors:** Ricardo Romero, Flávio S J de Souza

## Abstract

Many lizards (Squamata), as well as the tuatara (Rhynchocephalia), are distinguished among vertebrate groups for the presence of the parietal eye - also called “third eye” - a structure derived from the pineal complex that develops from the roof of the diencephalon and resembles a simplified retina. The parietal eye is located near the dorsal surface of the head and possesses photoreceptor cells expressing an array of nonvisual opsins that differs from the visual opsin repertoire of the lateral eyes. These pineal opsins are pinopsin (OPNP), parapinopsin (OPNPP) and parietopsin (OPNPT), all being evolutionary close to the visual opsins. A fourth member of the group, vertebrate-ancient opsin (OPNVA), is expressed in the brain. Here, we have searched over 50 lepidosaurian genomes (tuatara + lizards) for pineal non-visual opsins to check for the evolutionary trajectory of these genes during reptile evolution. Unexpectedly, we identified a novel opsin gene, which we termed “lepidopsin” (OPNLEP), that is present in the tuatara and most lizards but absent from the genomes of other reptiles. Phylogenetic analyses indicate that OPNLEP proteins are grouped in a clade distinct from nonvisual and visual opsins. Remnants of the gene are found in the coelacanth and some ray-finned fishes like gars and sturgeons, implying that OPNLEP is an ancient opsin that has been repeatedly lost during vertebrate evolution. As for the survey, we found that the tuatara and most lizards of the Iguania, Anguimorpha, Scincoidea and Lacertidae clades, which possess a parietal eye, harbour all five non-visual opsin genes analysed. Lizards missing the parietal eye, like geckos (Gekkota), the fossorial *Rhineura floridana* (Amphisbaenia) and lacertoids of the Teiidae and Gymnophthalmidae families lack most or all pineal nonvisual opsins. In summary, our survey of reptile pineal non-visual opsins has revealed i) the persistence of a previously unknown ancient opsin gene – OPNLEP - in lepidosaurians; ii) losses of non-visual opsins in specific lizard clades and iii) a correlation between the presence of a parietal eye and the genomic repertoire of pineal non-visual opsins.

## Introduction

In most animals, detecting light is an essential sensorial capability, employed to navigate and interact with the surrounding environment. In vertebrates, the best understood system of light detection is comprised by the two lateral eyes, which contain a layer of photoreceptor cells in the retina that absorbs photons and transmits visual information to the brain (Lamb 2013; Hagen et al. 2023). However, vertebrates can also detect light with other structures, like the pineal complex, a small group of sensory and endocrine brain organs derived from evaginations of the roof of the diencephalon during development. In mammals, the pineal complex is reduced to the pineal gland (epiphysis), which does not detect light by itself and serves a neuroendocrine function by secreting the hormone melatonin during nighttime. The pineal complex of other vertebrates, however, exhibits light sensitivity and may include other organs apart from the pineal proper (Oksche 1965; Dodt 1973; Eakin 1973; Vigh et al. 2002; Ekström and Meissl 2003; Mano and Fukada 2007). For instance, lampreys and teleost fishes have both a pineal and a parapineal organ with photoreceptor and neuroendocrine functions (Koyanagui et al, 2004; Koyanagui et al. 2015). In anuran amphibians, the pineal complex comprises a deep pineal and a frontal organ (Stirnorgan), located under the skin between the eyes and capable of detecting light directly (Adler 1976; Korf and Liesner 1981). The frontal organ and the deep pineal emerge from the same primordium during anuran development (van de Kamer et al. 1962).

The pineal complex of the tuatara and many lizards comprises a deep neuroendocrine pineal and a parietal eye, located on the dorsal part of the head, often visible on a head scale as a parietal “spot” (Gundy and Wurst. 1976a; Labra et al. 2010). The structure of the parietal eye resembles a lateral eye, with a lens and a simplified retina, being often nicknamed “third eye” for this reason, even though it cannot form images (Stebbins and Eakin 1958; Eakin 1973; Ung and Molteno 2004). In contrast to the anuran frontal organ, the evagination that gives rise to the reptilian parietal eye can be distinguished from the adjacent pineal evagination during development, indicating that it is likely homologous to the parapineal organ found in lampreys and teleosts (Oksche 1965; Eakin 1973; Meiniel 1977; Vigh et al. 2002). Anatomically, the parietal eye occupies an opening on the dorsal part of the skull – the pineal or parietal foramen – that allows exposure to light and the communication between the parietal eye and the brain through the parietal nerve (Engbretson et al. 1981). The presence of the parietal foramen in extant reptiles indicates the presence of a parietal eye and can be used to infer the presence of this sensory structure in extinct vertebrate lineages with ossified crania (Edinger 1955; Trost 1956; Benoit et al. 2016; Smith et al. 2018).

The retina of the parietal eye is lined with photoreceptor cells with a morphology similar to those of the lateral eyes, with a cilium-derived outer segment containing a dense arrangement of membranes (Eakin 1973; Solessio and Engbretson 1993). The capacity of detecting light depends on the presence of opsin proteins embedded in membrane stacks within the outer segments of the photoreceptors. Opsins are G protein-coupled transmembrane receptors responsible for detecting light. Their photosensitivity depends on a small chromophore moiety, like 11-*cis*-retinal, which is covalently bound to the opsin protein and can absorb photons. Light converts 11-*cis*-retinal to all-*trans*-retinal, triggering a conformational change of the opsin molecule and the activation of a G protein (Terakita 2005; Terakita et al. 2012). The ancestral, basic repertoire of vertebrate visual opsins comprises five members, namely RHO/RH1, RH2, LSW, SWS1 and SWS2, each one with a particular light absorption profile. Vertebrate genomes, however, harbour many other opsin genes, which can be grouped into five families (OPN1, OPN3, OPN4, OPN5 and OPN6) following their phylogenetic relationships (Beaudry et al. 2017; Pérez et al. 2018).

Vertebrate visual opsins are included into the OPN1 group together with four so-called “nonvisual opsins”, namely i) pinopsin (OPNP), ii) parapinopsin (OPNPP), iii) parietopsin (OPNPT) and iv) vertebrate-ancient opsin (OPNVA). Many of these are expressed in photoreceptor cells of the pineal complex of non-mammalian vertebrates (Kawano-Yamashita et al, 2014; Pérez et al, 2018). OPNP has an absorption profile in the blue range (∼470 nm; Okano et al, 1994; Sato et al, 2018) and its expression has been detected in the pineal gland of several vertebrates like the chick, lamprey, the frog, *Xenopus laevis* and some lizards (Okano et al. 1994; Max et al. 1995; Kawamura and Yokoyama 1997; Yokoyama and Zhang 1997; Frigato et al. 2006; Bertolesi et al. 2020) as well as the retina of a gecko lizard and several non-teleost fishes (Taniguchi et al. 2001; Sato et al. 2018). In the parietal eye of lizards, OPNP protein expression is found in the outer segments of photoreceptors of the iguanid *Uta stansburiana* together with OPNPT (Su et al. 2006). OPNPP has an absorption maximum in the ultraviolet range (∼370 nm; Koyanagi et al. 2004) and is expressed in the pineal complex of teleost fishes and lamprey (Blackshaw and Snyder 1997; Koyanagi et al. 2004, 2015), and the frog *X. laevis* (Bertolesi et al. 2020). In the parietal eye, OPNPP is coexpressed with OPNPT in photoreceptors of *Iguana iguana* (Wada et al. 2012). OPNPT, which absorbs light in the green range (∼522 nm; Su et al. 2006; Sakai et al, 2012) is expressed in the pineal complex of the zebrafish and *X. laevis* (Wada et al. 2018; Bertolesi et al. 2020). As mentioned, OPNPT protein is detected in the outer segments of photoreceptors of the parietal eye of iguanids, being coexpressed with OPNP in *U. stansburiana* and with OPNPP in *I. iguana* (Su et al. 2006; Wada et al. 2012). OPNVA, with a ∼500 nm absorption maximum (Sato et al. 2011), is mostly expressed in cells located in the deep brain of vertebrates and its expression in the pineal complex has not been detected, except in the pineal of the Atlantic salmon (Kojima et al. 2000; Philp et al. 2000; Bertolesi et al. 2020).

Although only a few species have been studied, previous work indicates that the parietal eyes of lizards express OPNP, OPNPP and OPNPT. The photoreceptors of the parietal eye have a very unusual behaviour in that they display chromatic antagonism: they depolarise in response to green light and hyperpolarise in response to blue light (Solessio and Engbretson 1993). Studies on *U. stansburiana* indicate that this antagonistic behaviour is mediated by nonvisual opsins expressed together in the same photoreceptors (Su et al. 2006). Thus, the green light-responsive OPNP activates the G protein gustducin, leading to the activation of a cGMP-phosphodiesterase, a decrease in cGMP levels, the closure of cyclic nucleotide-gated channels (CNG) and the depolarisation of the photoreceptor. The blue light-responsive OPNPT, on the other hand, activates Go, leading to the inhibition of a cGMP-phosphodiesterase, an increase in cGMP, the opening of CNGs and the hyperpolarisation of photoreceptor cells (Su et al. 2006; Kawano-Yamashita et al. 2014). Interestingly, the colocalisation of OPNPP and OPNPT in photoreceptors of *I. iguana* indicates that a similar antagonistic chromatic response might exist between UV and green light detected by OPNPP and OPNPT, respectively (Wada et al. 2012). The photoreceptors synapse with ganglion cells located in the outer layer of the parietal eye retina, which project to the dorsolateral portion of the left habenula and other brain areas (Engbretson et al. 1981; Engbretson 1992).

The functions of the parietal eye on lizard behaviour and physiology have been studied in several species, usually by experiments in which the parietal eye is removed or covered to prevent it from detecting light (Tosini, 1997). Lizards are environmental thermoregulators, and blocking parietal eye function alters the duration of exposure to sunlight and body temperature, as well as daily activity patterns (Stebbins and Eakin 1958; Huntchinson and Kosh 1974; Tosini 1997; Traeholt 1997). There is also evidence that the parietal eye is necessary for lizards to orient themselves in space (Ellis-Quinn and Simony 1991; Freake 2001) likely by using polarised sunlight as a compass (Beltrami et al. 2010). It may also be required for the control of metabolic rate and reproduction, possibly by regulating melatonin production by the pineal organ (Tosini 1997). In spite of the important functions associated with it, the parietal eye is not present in all lizards. Gundy and Wurst (1976a) found that 12 out of 18 families of “Larcetilia” (excluding amphisbaenians and snakes) had parietal eyes, with all members of some important families, like Gekkota, Dibamidae and Teiidae lacking parietal eyes altogether. In addition, snakes, which are phylogenetically grouped within Anguimorpha and Iguania in the Toxicofera clade, also lack parietal eyes.

Even though nonvisual OPN1 opsins are ancient genes, previous studies have found that these genes have a somewhat patchy distribution in vertebrate groups. In particular, *OPNP, OPNPP* and *OPNPT*, which are expressed in the lizard parietal eye, have frequently been lost during evolution. Thus, snakes have lost *OPNP, OPNPP* and *OPNPT*, turtles have lost *OPNPP* and *OPNPT* and crocodylians, which completely lack a pineal complex, lost *OPNP, OPNPP* and *OPNPT* (Emerling 2017a, 2017b). Birds lack *OPNPP* and *OPNPT*, while mammals are unique in having lost all nonvisual OPN1 opsins (Hankins et al. 2014; Emerling 2017a). In contrast, the genome of frogs *X. laevis* and *X. tropicalis*, which have a frontal organ similar to the parietal eye, contain all nonvisual OPN1 opsins (Bertolesi et al. 2020). The presence of these opsins seems to be linked to the occurrence of photosensitive organs of the pineal complex of vertebrates, like the anuran frontal organ and the reptilian parietal eye. Indeed, *OPNP, OPNPP* and *OPNPT* are all expressed in the developing pineal complex of *X. laevis* (Bertolesi et al. 2020).

Lizards and snakes comprise the Squamata, with over 11,000 species (including over 7,600 lizards) displaying a high diversity of form and behaviour (Meiri 2024). Together with Rhynchocephalia, of which the tuatara of New Zealand (*Sphenodon punctatus*) is the sole extant representative, Squamata is included within Lepidosauria. It is estimated that Rhynchocephalia and Squamata diverged in the late Permian, before 250 million years ago (MYA), while extant squamate groups diversified during the Jurassic and later (after 200 MYA, Simões et al. 2020, 2022). To extend the analyses of pineal photosensitivity and opsin evolution to this understudied group of animals, we sought in this work to identify the repertoire of genes encoding nonvisual OPN1 opsins in the genomes of Lepidosauria. In this process, we discovered a new nonvisual OPN1 opsin, *lepidopsin* (*OPNLEP*), which is phylogenetically similar to *OPNPP* and *OPNPT*. In lizard groups, we found that the presence of a parietal eye is associated with the presence of *OPNPP, OPNPT* and *OPNLEP*, while *OPNP* is frequently lost. Fragments of *OPNLEP* exons are seen in the genomes of snakes, turtles and crocodylians, showing that this gene was once functional in all stem reptilians. Surprisingly, fragments of the *OPNLEP* gene are present in the genome of the coelacanth and some basal groups of ray-finned fishes, implying that the gene was repeatedly lost during evolution, likely due to the loss of the parietal eye in most sarcopterygian and tetrapod clades.

## Results

### Discovery of a new opsin: OPNLEP

A search for nonvisual opsins of the OPN1 group in lizard genomes readly reveals the presence of all four known genes, namely *OPNP, OPNPP, OPNPT* and *OPNVA* (see below). Unexpectedly, we found a fifth nonvisual OPN1 opsin gene in the tuatara and in most lizard genomes, which we named ***Lepidopsin*** (***OPNLEP***) for its occurrence in Lepidosauria. The predicted aminoacid sequence of OPNLEP proteins ranges from 376 to 385 residues and it aligns well with that of other opsins, including the seven transmembrane (TM) domains (Fig. 1a and supplemental fig. S1). It has all the typical conserved residues found in opsins, like cysteines 110 and 185, that form a disulfide bond, and lysine 296 in TM7, which binds the retinal moiety necessary for light absorption (Fig. 1b, residues are numbered according to bovine rhodopsin). Interestingly, glutamate 113, that serves as counterion to the protonated Schiff base of the retinal chromophore in most OPN1 opsins, is changed to glutamine in OPNLEP, as in OPNPT (Su et al, 2006; Fig. 1b). Thus, it is probable that the Schiff base counterion in OPNLEP is glutamate 181, as has been shown for OPNPT (Sakai et al. 2012).

**FIG. 1.**
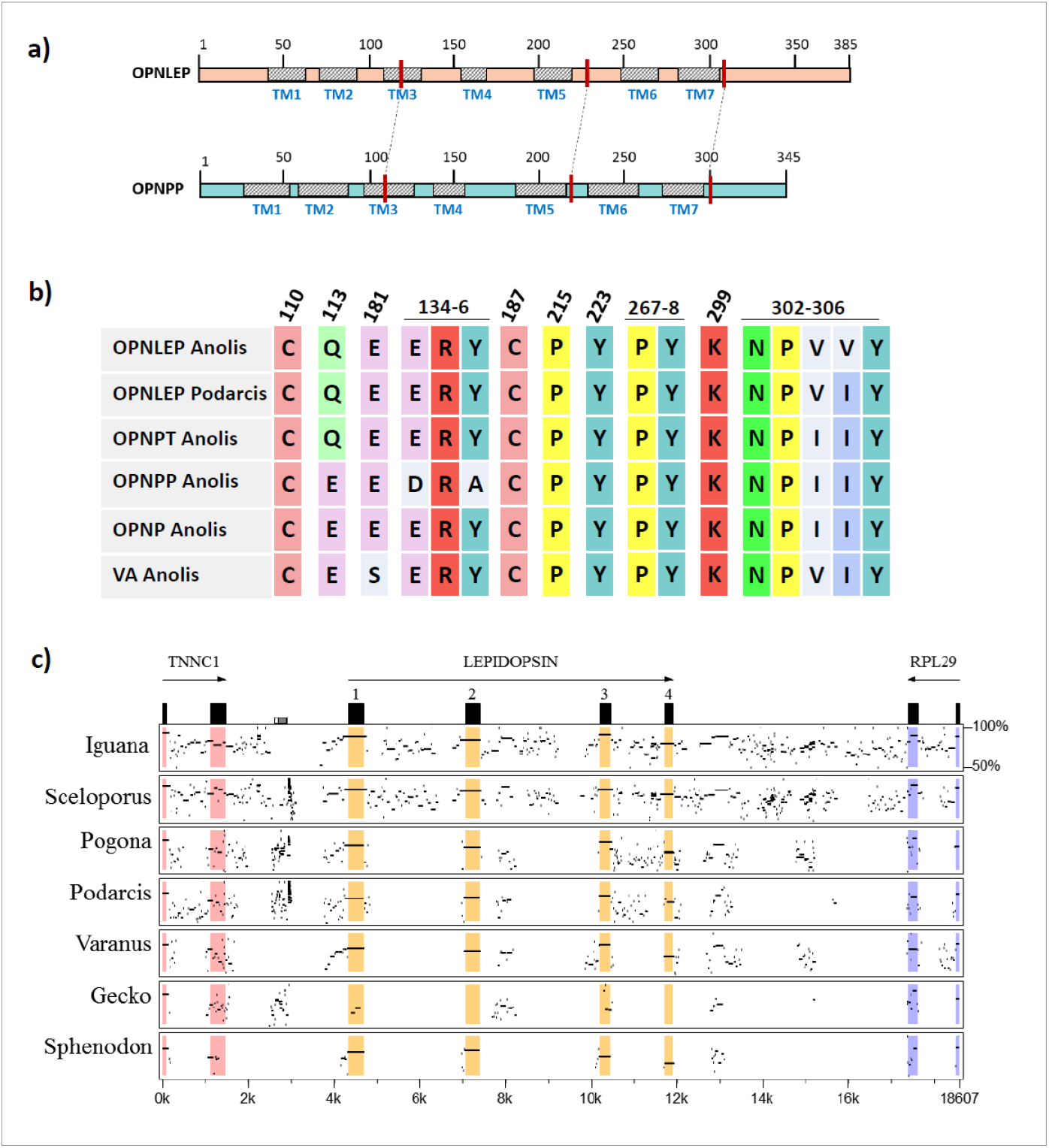
Characterisation of a new reptilian opsin, lepidopsin. a) Schematics of the structure of OPNLEP and OPNPP proteins of the green anole, *A. carolinensis*. TM: transmembrane domains 1-7. Red bars and dashed lines point to the approximate positions of the three introns in both genes. b) Key residues involved in opsin function in five nonvisual OPN1 opsins from the green anole and the OPNLEP of the common wall lizard (*Podarcis muralis*). Residue numbers correspond to bovine rhodopsin. c) Percentage identity plot (PipMaker program) between the nucleotide sequence of the green anole *OPNLEP* locus and homologous loci from other lepidosaurian species: *Iguana delicatissima, Sceloporus undulatus, P. muralis, Varanus komodoensis, Gekko japonicum* and *S. punctatus*. The last two exons of *TNNC1* and *RPL29* are also shown in the plot.

In all lepidosaurians analysed, the *OPNLEP* gene is flanked by *TNNC1* (encoding troponin c1) and *RPL29* (encoding ribosomal protein L29; Fig. 1c). A global nucleotide sequence alignment analysis with MultiPipMaker (Schwartz et al. 2000) using as reference the *OPNLEP* locus of the green anole (*Anolis carolinensis*) shows that the four exons of the gene are highly conserved in lepidosaurians (Fig. 1c). The exons of the green anole are particularly evident in the alignment with the tuatara locus, confirming that *OPNLEP* has four coding exons. This is the same number of exons as *OPNPP* and *OPNPT*, while *OPNP* and *OPNVA* genes have five. In addition, the four predicted exons of *OPNLEP* correspond to the same, homologous exons of *OPNPP* and *OPNPT* (Fig. 1a). Interestingly, in species with chromosome-level sequences, we found *OPNLEP* and *OPNPP* are located in the same chromosomes separated by similar distances, while other nonvisual OPN1 genes are found in different chromosomes (supplemental table S1). This suggests that *OPNLEP* and *OPNPP* may have originated by a segmental duplication. A further analysis of the origin of *OPNLEP* is given in a section below.

### Survey of nonvisual OPN1 opsins in lepidosauria

We took advantage of the availability of the tuatara and many lizard genomes to search for nonvisual opsins of the OPN1 group, including OPNLEP (see Methods). As a general evolutionary framework, we followed recent molecular phylogenetic analyses of squamates (Pyron et al, 2013; Burbrink et al, 2020; Simões and Pyron, 2021). Our survey includes representatives of Rhynchocephalia (the tuatara) and most large clades of lizards, namely Gekkota (8 species from 4 families), Iguania lizards belonging to Pleurodonta (10 species from 4 families) and Acrodonta (8 species from 2 families), Lacertoidea (14 species from 4 families), Anguimorpha (6 species from 5 families) and Scincoidea (7 species from 2 families). Although included in Squamata, we have not analysed snake (Serpentes) genomes as previous analyses have already shown the loss of *OPNP, OPNPP* and *OPNPT* genes from this clade (Emerling, 2017b), and we could not find the complete *OPNLEP* gene in snakes (see below).

OPNP, OPNPP and OPNPP are all known to be expressed in the parietal eye of lizards (Kawamura and Yokoyama 1997; Taniguchi et al. 2001; Su et al. 2006; Wada et al. 2012), but this structure is not present in all groups. To check for a correlation between opsin repertoire and the presence of the parietal eye, we considered the occurrence of a parietal eye or a parietal “spot” from data gathered by Gundy and Wurst (1976a, 1976b) and Lee (1998), as well as by the presence of a pineal opening or parietal foramen in the skull, which generally indicates the presence of a parietal eye (Edinger 1955; Trost 1956; see Materials & Methods).

The distribution of the OPN1 nonvisual opsins in 54 lepidosaurian genomes is shown in Figure 2. Our observations can be summarised as follows:

**FIG. 2.**
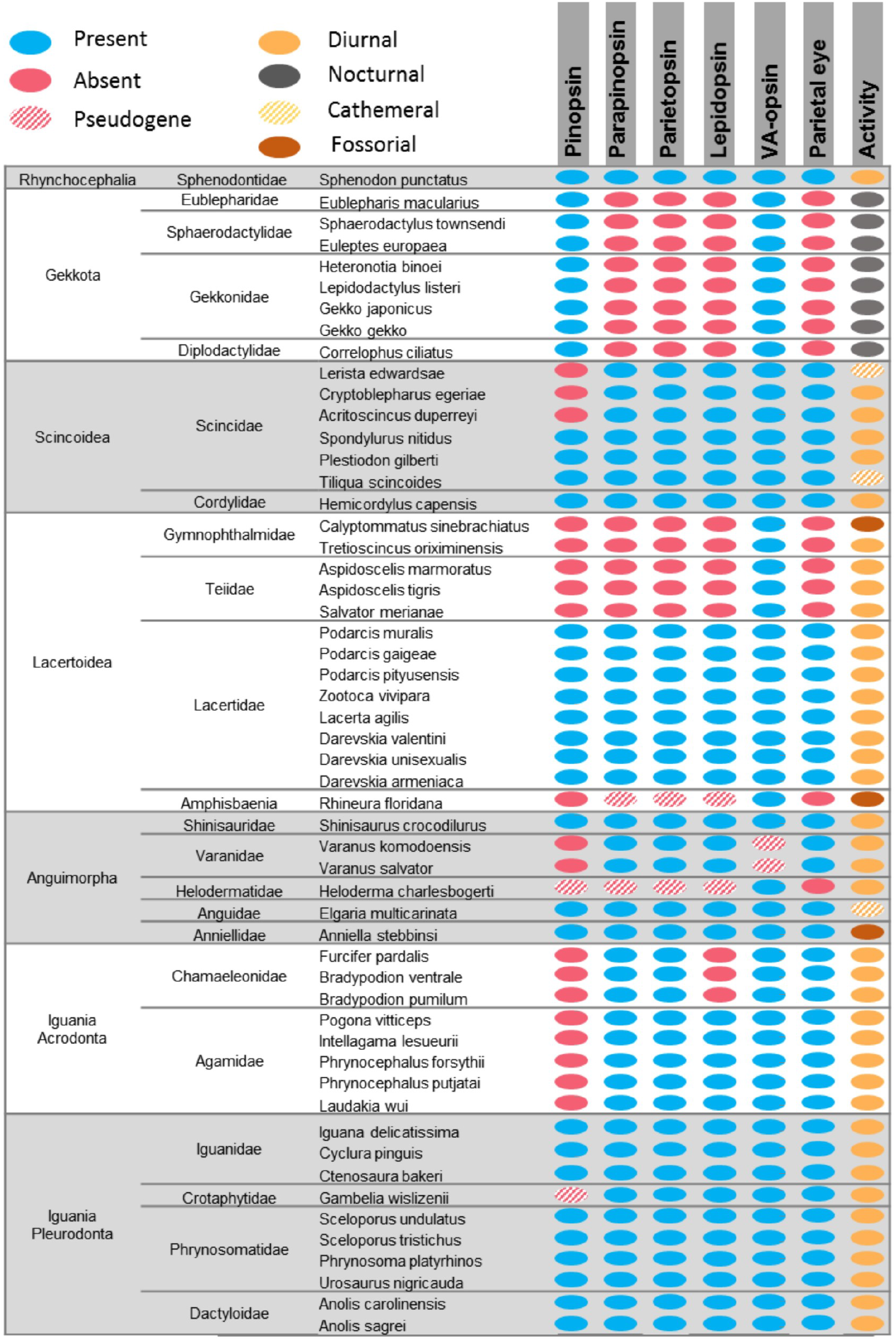
Distribution of nonvisual OPN1 opsins among lepidosaurians. The presence of a parietal eye, as well as patterns of activity or fossoriality, are indicated.

#### Rhynchocephalia

the tuatara genome has all opsins surveyed, including *OPNLEP*. The tuatara has a well-developed parietal eye, with a retina and lens, placed inside a large parietal foramen (Dendy 1911; Edinger 1955; Ung and Molteno 2004).

#### Gekkota

gecko genomes have intact *OPNP* and *OPNVA* genes, but all species surveyed have lost the other three opsin genes. Geckos are known to have lost the parietal eye (Gundy and Wurst 1976a, 1976b). In contrast to other lizards, most gecko species are nocturnal and may have experienced a “nocturnal bottleneck” during their early evolution (Pinto et al. 2019; Katti et al. 2019; Kojima et al. 2021).

#### Scincoidea

the African girdle lizard *Hemicordylus capensis* (Cordylidae) has all opsins surveyed. As for skinks (Scincidea) all have all opsins except for *OPNP*, which is lost in three of the species we analysed. Both cordylids and skinks have parietal eyes (Gundy and Wurst 1976b).

#### Lacertoidea

lizards of the Lacertidae family have all five opsins and well-developed parietal eyes (Trost 1956; Gundy and Wurst 1976a, 1976b). The opsin repertoire and status of the parietal eye in other families are different. Teiids and gymnophthalmids are related groups (Goicoechea et al. 2016) that lack a parietal eye (Gundy and Wurst 1976b; Lee 1998). Interestingly, the four species surveyed lack all opsins except for *OPNVA*. For amphisbaenians, which lack a parietal eye and are of fossorial habits (Lee 1998), we analysed the *Rhineura floridana* genome (Florida worm lizard). *R. floridana* has recognisable *OPNPP, OPNPT* and *OPNLEP* genes that nevertheless carry mutations that should render them pseudogenes (supplemental table S2). It lacks *OPNP* and its only nonvisual OPN1 gene that seems to be funcional is *OPNVA*.

#### Anguimorpha

monitor lizards of the Varanidae family have a well-developed parietal eye, and the two species that we surveyed (*Varanus komodoensis* and *V. salvator*) have *OPNPP, OPNPT* and *OPNLEP*, while *OPNP* is lost and the *OPNVA* gene lacks its first and fifth exons and is thus pseudogenised. The representatives of Anguidae and Shinisauridae families, on the other hand, have all five opsin genes and also possess a parietal eye. *Anniella stebbinsi* also has all opsin genes surveyed, even though it belongs to a small group of Californian legless lizards of fossorial habit (Anniellidae; Papenfuss and Parham 2013). Interestingly, *Anniella pulchra*, a related species, lacks a parietal foramen but still has a well-formed parietal eye located under the skull (Gundy and Wurst. 1976a). Finally, the Guatemalan beaded lizard (*Heloderma charlesbogerti*), belonging to a family (Helodermatidae) lacking parietal eyes, has all nonvisual OPN1 opsins in a pseudogenised state except for *OPNVA*.

#### Pleurodont iguanians

pleurodonts have all five opsins, although one species, *Ctenosaura bakeri*, has a pseudogenised *OPNP*. Iguanians have a well-developed parietal eye (Trost 1956; Gundy and Wurst 1976a, 1976b), and indeed the expression of *OPNP, OPNPT* and *OPNPP* expression has been studied in detail in the pineal eye photoreceptors of two pleurodont species, namely *Uta stansburiana* and *Iguana iguana* (Su et al. 2006; Wada et al. 2012).

#### Acrodont iguanians

both agamids and chamaeleonids lack *OPNP*, while chamaleonids lack, in addition, *OPNLEP*. Both groups, however, have parietal eyes (Trost 1956; Gundy and Wurst 1976a, 1976b). Interestingly though, chamaeleonids of the *Chameleo* genus have a somewhat degenerate parietal eye, described as a hollow vesicle lacking lens and retina (Edinger 1955; Gundy and Wurst 1976a).

In general, we observe that genomes from lepidosaur species that have parietal eyes possess *OPNPP, OPNPT* and *OPNLEP*, while species and groups where this structure is missing (geckos, teiids, gymnophthalmids, amphisbaenians, helodermatids) lack all three functional genes.

### Evolutionary origin of OPNLEP

A complete gene for the new nonvisual OPN1 opsin, *OPNLEP*, is found in 36 out of 54 lepidosaurian species, and 12 out of 22 lepidosaurian families, that we analysed (Fig. 2). In a phylogenetic analysis of lepidosaurian OPN1 opsin proteins, including representatives of the five basic visual opsins (LSW, SWS1, SWS2, RH1 and RH2), OPNLEP sequences are grouped together in a well-defined branch with high statistical support (Fig. 3). OPNP clusters with visual opsins, as observed in previous studies (Beaudry et al. 2017; Bertolesi et al. 2020) while OPNLEP is found closest to OPNPP and OPNPT. A close-up of the OPNLEP branch shows the expected topology, with the tuatara sequence at the base of the clade and the other branches corresponding to the lizard taxonomic groups analysed (supplemental fig. S2).

**FIG. 3.**
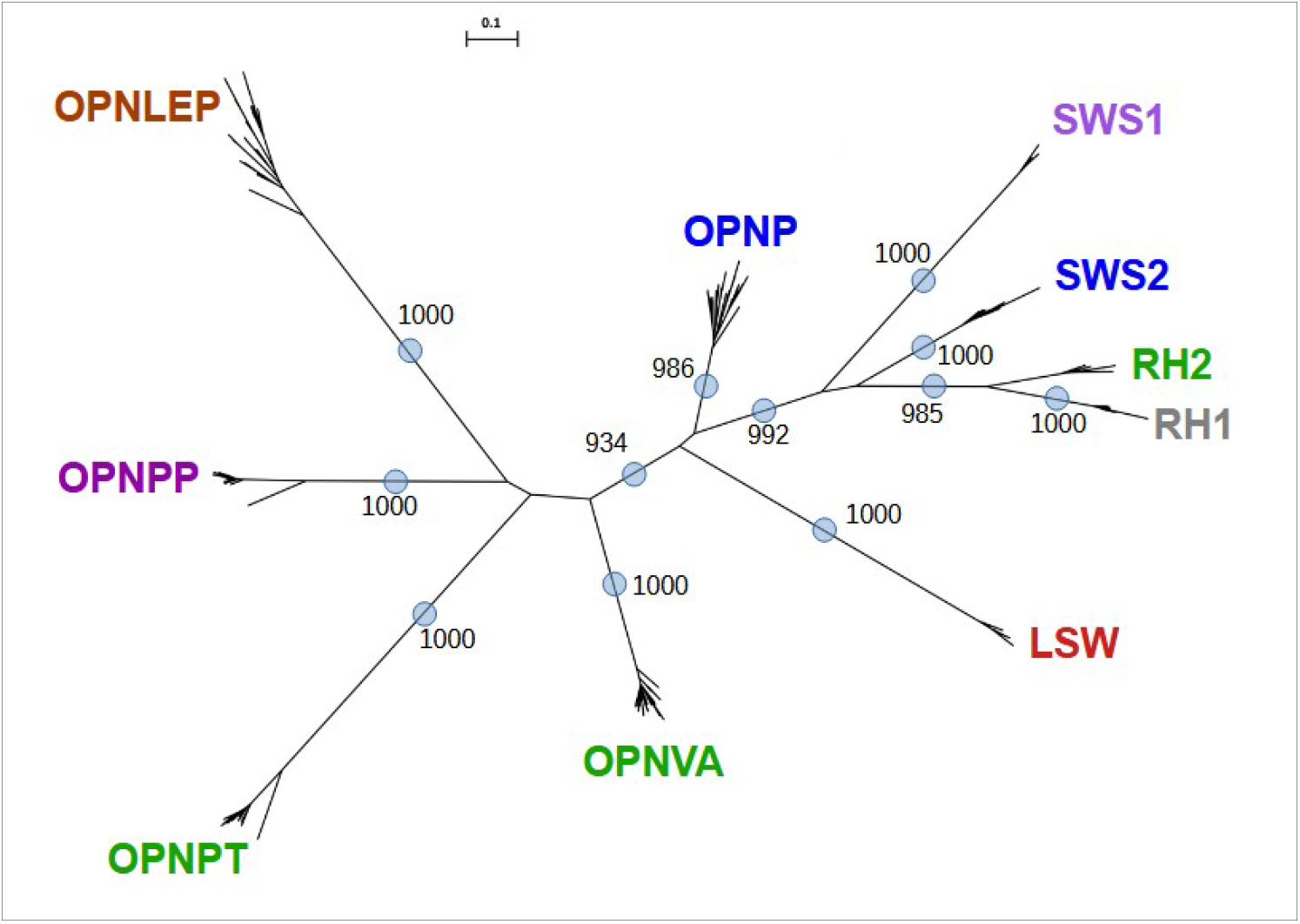
OPNLEP proteins comprise a new nonvisual opsin clade. Unrooted phylogenetic tree of nonvisual OPN1 opsin proteins and a selection of visual opsins of lepidosaurians estimated using the balanced minimum evolution method (FastME 2.0). Label colours indicate approximate absorption maximum of opsins, except for OPNLEP which is currently unknown. Individual species and proteins are not shown for clarity. Blue circles indicate branches supported by a bootstrap >900 out of 1000 replicates. Tree scale is shown.

Many species of lizards have a *OPNLEP* gene that carries several mutations that render them pseudogenes, as is the case of the amphisbaenian *R. floridana* and the beaded lizard, *H. charlesbogerti* (supplemental table S2). In geckos, fragments of the *OPNLEP* gene are still present, but not the whole gene. For instance, the nucleotide global alignment of Fig. 1c shows that fragments of exon 1 and 3 of *OPNLEP* are still detectable in *Gekko japonicus*, while other exons are absent.

Although BLAST searches indicate that a complete *OPNLEP* gene is absent from other reptiles, we used MultiPipMaker to check for remnants of the gene between *TNNC1* and *RPL29* in the genomes of other reptile clades. We found that fragments of *OPNLEP* exons are recognisable in snakes like *Candoia aspera* and *Thamnophis elegans*, turtles like *Chelonia mydas* and *Gopherus evgoodei* and crocodilians like *Alligator mississippiensis*, indicating that the origin of *OPNLEP* predates lepidosaurians (Fig. 4 and supplemental table S3).

**FIG. 4.**
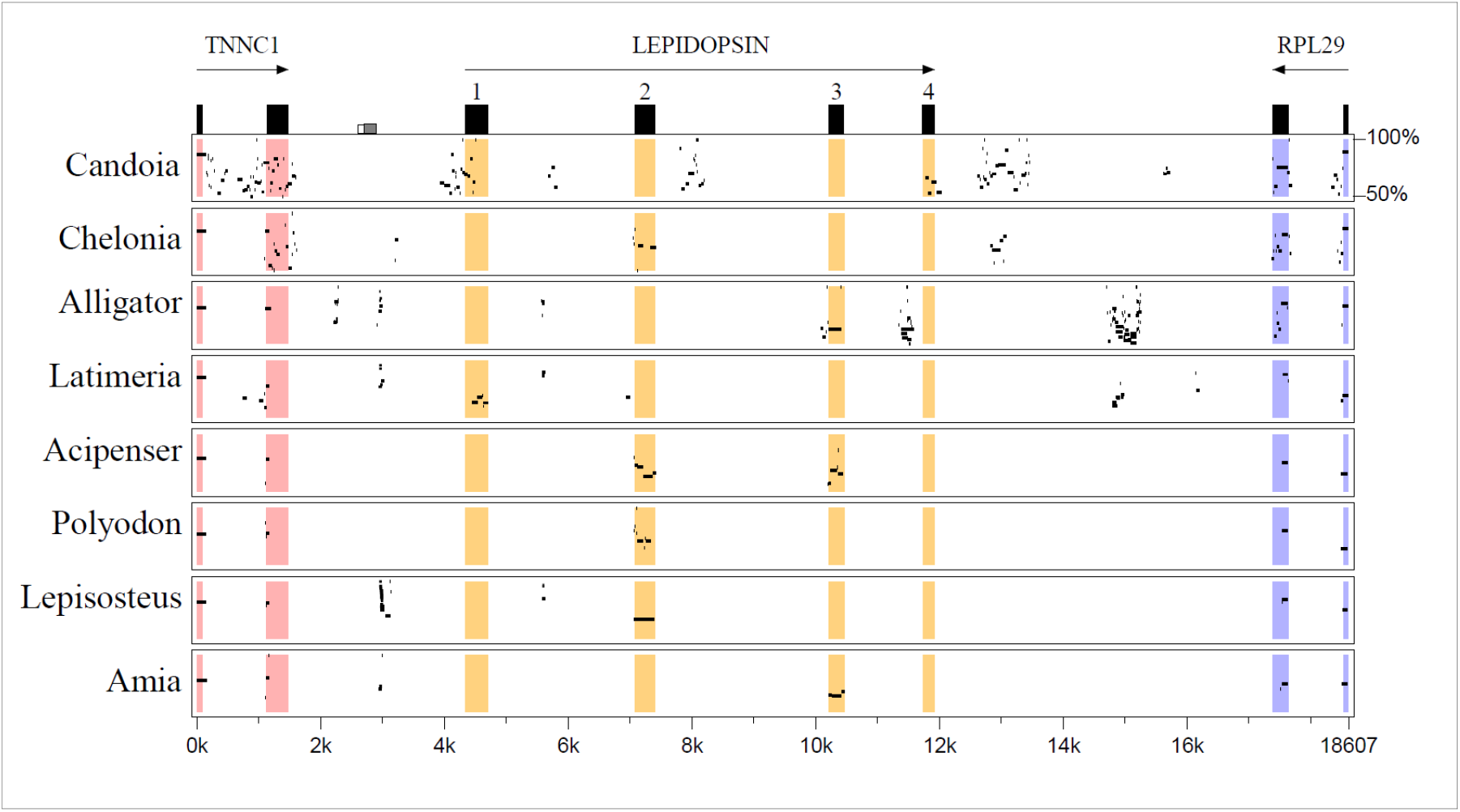
Remnants of *OPNLEP* exons are still found in several vertebrate clades. Percentage Identity Plot (MultiPipMaker) of the green anole *OPNLEP* locus and the homologous loci of the Papuan ground boa (*Candoia aspera*), green sea turtle (*Chelonia mydas*), alligator (*Alligator mississipiensis*) and coelacanth (*Latimeria chalumnae*). Ray-finned fishes represented are the sterlet (*Aciperser ruthenus*), paddlefish (*Polyodon spathula*), spotted gar (*Lepisosteus oculatus*) and bowfin (*Amia calva*). The two last exons of both flanking genes, *TNNC1* and *RPL29*, are present in all species.

When we extended the search to other lobe-finned fishes (Sarcopterygii), we were unable to identify *OPNLEP* remnants in amphibians, birds and mammals, as well as lungfishes (supplemental fig. S3). Remarkably, most of the exon 1 of *OPNLEP* is readily recognisable in the genome of the coelacanth, *Latimeria chalumnae*, between the homologues of *TNNC1* and *RPL29* (Fig. 4). Among ray-finned fishes (Actinopterygii), long remnants of *OPNLEP* exons were detectable in the genomes of members of the Holostei clade, namely the spotted and longnose gars (*Lepisosteus oculatus* and *L. osseus*) and the bowfin (*Amia calva*), as well as Acipenseriformes like the sturgeon (*Acipenser ruthenus*) and paddlefish (*Polyodon spathula*), in all cases between the *TNNC1* and *RPL29* genes (Fig. 4 and supplemental table S3). *OPNLEP* was not retrieved from the genomes of bichirs (Polypteriformes) and teleosts (Teleostei). Among other vertebrate groups, we were unable to detect *OPNLEP* remnants in cartilaginous (Chondrichthyes) or jawless (Cyclostomata) fishes (supplemental table S3). These results indicate that *OPNLEP* is an ancient opsin gene that originated before the divergence of lobe- and ray-finned fishes but has been retained only in recent lepidosaurians.

## Discussion

In this work, we describe the repertoire of nonvisual opsins of the OPN1 group in available lepidosaurian genomes. Our main observations are that **i)** lepidosaurians have a new OPN1 gene, *OPNLEP*, most similar to *OPNPP* and *OPNPT*; **ii)** the repertoire of nonvisual opsins varies among lizards, with a tendency for lizard taxa lacking the parietal eye also lacking *OPNPP, OPNPT* and *OPNLEP* genes; **iii)** *OPNLEP* is an ancient gene, having originated at least in the begining of the evolution of bony fishes.

In a phylogenetic tree, OPNLEP proteins appear closer to OPNPP and OPNPT than to other OPN1 opsins. The three genes have four exons and three introns with splicing sites located in the same relative positions, indicating an origin by duplication from an ancestral gene. In addition, *OPNLEP* and *OPNPP* genes are always linked in the same chromosome of lizards. This observation aligns with the scenario proposed by Lagman et al (2024), in which several rounds of tandem duplications gave origin to OPN1 opsin genes in ancient vertebrates. In fact, the gene referred by them as “parapinopsin-like” in the spotted gar genome (Langman et al. 2024) might correspond to the *OPNLEP* fragment in this species. The aminoacid sequence of OPNLEP resembles OPNPT in having a glutamine at position 113, instead of glutamate, so it is likely that the conserved glutamate 181 in OPNLEP is the counterion to the Schiff base linkage between the retinal chromophore and lysine 299, as is the case for OPNPT and invertebrate opsins (Sakai et al. 2012). The absorption spectrum of OPNLEP, as well as its characterisation as a monostable or bistable pigment, awaits photochemical studies with reconstructed recombinant protein.

Our survey of nonvisual OPN1 opsins in lepidosaurians revealed some important tendencies in the evolution of these genes, summarised in Figure 5a. The tuatara has maintained the ancestral repertoire of five nonvisual opsins. In lizards, *OPNVA* seems to be the least likely to be lost, missing only from varanids. *OPNVA* is expressed in neurons located in the deep brain, presumably rendering it less dependent on the function of the eyes and the pineal complex. It is also present in snakes (Emerling 2017b). *OPNP*, on the other hand, was lost in several clades, including all agamids, several skinks, varanids, teiids and gymnophthalmids, as well as the beaded lizard and the amphisbaenian, *R. floridana*. It is expressed in the retina and parietal eyes of lizards (Taniguchi et al. 2001; Su et al. 2006; Kato et al. 2018). *OPNPP, OPNPT* and *OPNLEP* are present in lizards exhibiting a parietal eye and missing in clades where this sensory structure is not present. Thus, the three genes have been lost in geckos, teiids, gymnophthalmids, the beaded lizard and the fossorial amphisbaenian *R. floridana*, all of which lack parietal eyes. The only exception are chamaleonids, which have lost *OPNLEP* but maintained the other two opsins. The parietal eye of chameleons, at least those of the *Chameleo* genus, has a degenerated structure (“a hollow vesicle”, Gundy and Wurst 1976a), and it may not be as functional as in other agamids. Thus, in general, the conservation of *OPNPP, OPNPT* and *OPNLEP* in lepidosaurian genomes seems to depend on the presence of the parietal eye. Interestingly, this correlation is also observed in *Anniella stebbinsi*, which although being from a fossorial lizard family (Anniellidae) that lacks a parietal foramen, has a well-developed parietal eye under the skull (Grundy and Wurst 1976a, 1976b) and intact *OPNPP, OPNPT* and *OPNLEP* genes.

**FIG. 5.**
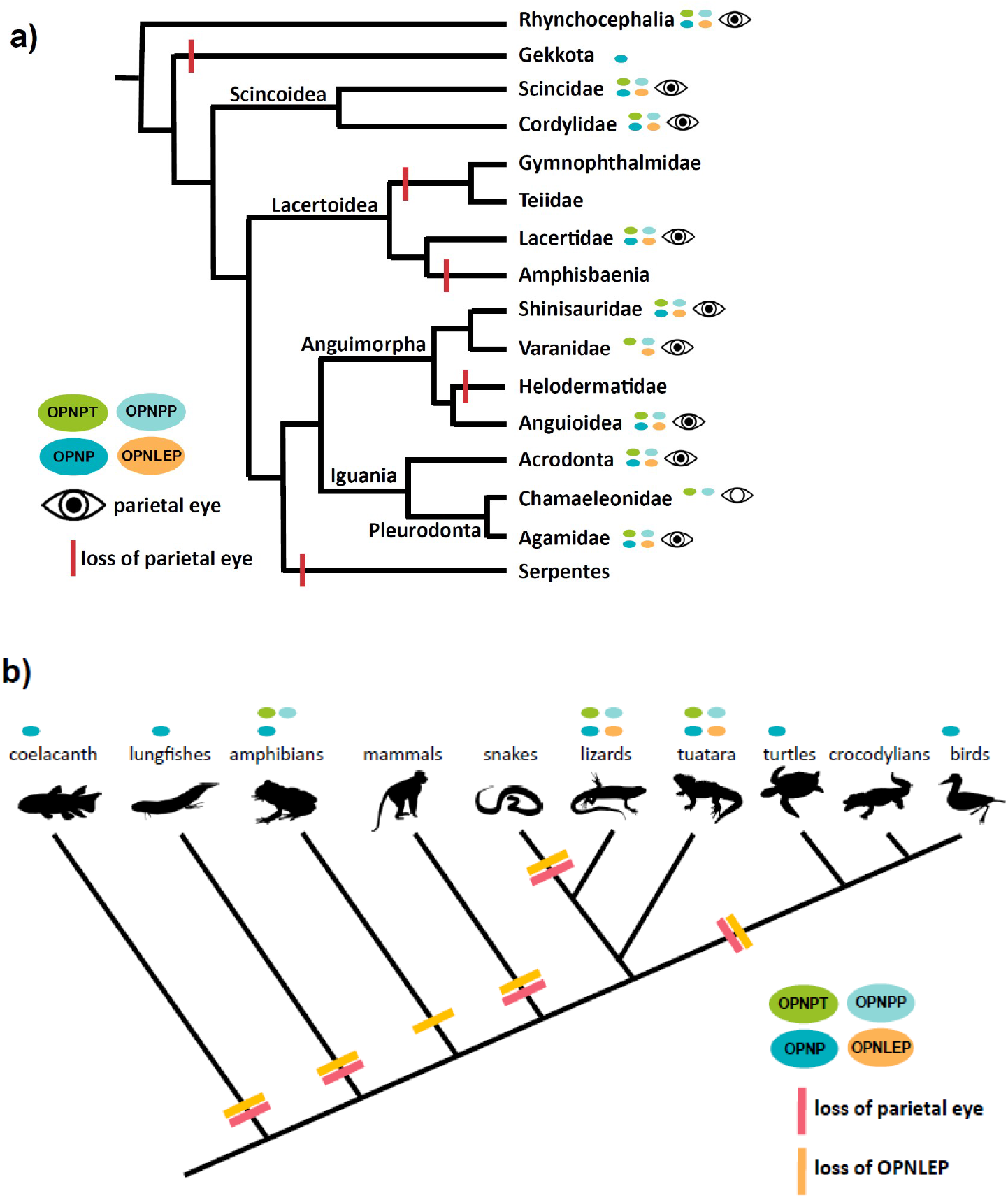
Evolution of OPN1 nonvisual opsins and parietal eye in vertebrates. a) Phylogenetic relationships between lepidosaurian clades showing the correlation between the presence of a parietal eye and pineal nonvisual opsins. OPNPP, OPNPT and OPNLEP tend to be present in clades with well-developed parietal eyes. b) Sarcoptergygian phylogenetic relationships and their repertoire of OPN1 nonvisual opsins. The parietal eye and OPNLEP have been independently lost several times during evolution.

In general terms, the fact that lizards with parietal eyes tend to have a complete repertoire of nonvisual OPN1 opsins is consistent with the observation that *OPNP, OPNPP* and *OPNPT* expression has been observed in photoreceptor cells of the lizard parietal eye (Su et al. 2006; Wada et al. 2012). *OPNP, OPNPP* and *OPNPT* are also expressed in the pineal complex of the frog *Xenopus laevis* (Bertolesi et al. 2020), which has a frontal organ similar to a parietal eye (van de Kamer et al. 1962; Eakin 1973). Thus, we consider very likely that future studies will reveal that *OPNLEP* is also expressed in the parietal eyes of lepidosaurians.

From an evolutionary and ecological standpoint, the profile of opsin losses in lepidosaurians generally agrees with the idea that nocturnality and fossoriality favour the reduction or loss of photosensitive organs, like the parietal eye, and associated genes. Snakes, which belong to the clade Toxicofera together with Iguania and Anguimorpha, seem to have evolved from nocturnal and/or fossorial ancestors (Hsiang et al. 2015), influencing adaptations like the loss of limbs, structural changes in retinal photoreceptors and loss of two visual opsins (*SWS2* and *RH2*; Simões et al. 2015; Emerling 2017b; Katti et al. 2019). Snakes have lost the parietal eye as well as *OPNP, OPNPP, OPNPT* and only remnants of the *OPNLEP* gene are still detectable. The gecko lineage also seems to have undergone a “nocturnal bottleneck” during evolution, when their eyes adapted to reduced light conditions and many genes were lost, including two visual opsins (*SWS2* and *RH1*; Emerling 2017b; Pinto et al. 2019; Katti et al. 2019; Kojima et al. 2021). Emerling (2017b) showed that *G. japonicus* lost *OPNPP* and *OPNPT*, and we have now extended this observation to *OPNLEP* and eight species of geckos from four different families, which suggests that the noctural lifestyle led to the loss of the parietal eye and these three opsins from this lineage. Amphisbaenians are legless fossorial lizards with thick scales, reduced eyes and missing parietal eyes. Consistent with this, *R. floridana* has lost *OPNP* and pseudogenised *OPNPP, OPNPT* and *OPNLEP*. It remains to be seen if the loss of these opsins will be a general feature of amphisbaenians. Finally, extant archelosaurians (turtles, crocodilians and birds) also lack parietal eyes and lost *OPNPP, OPNPT* (Emerling 2017a) and only have remnants of *OPNLEP*.

Remarkably, even though *ONPLEP* could not be retrieved from the genomes of amphibians, birds, mammals and other vertebrates, we did identify exon remnants in the coelacanth, the Australian lungfish and in two groups of ray-finned fishes, Acipenseriformes and Holostei. In all these species, the *OPNLEP* fragments are located in the same, homologous locus where the gene is found in lepidosaurians. The slow pace of molecular and morphological evolution of coelacanths (Amemyia et al. 2013; Cavin and Guinot 2014; Clement et al. 2024), sturgeons, the paddlefish, gars and bowfins (Redmond et al. 2023; Brownstein et al. 2024) may account for this survival. Thus, even though complete *OPNLEP* genes are only present in lepidosaurs, this opsin is an ancient one, dating from before the divergence between ray- and lobe-finned fishes, in the middle Silurian, around 430 million years ago (MYA; Brazeau and Friedman 2015). Among sarcopterygians, this implies that *OPNLEP* was independently pseudogenised and lost in the lineages leading to extant coelacanths, lungfishes, amphibians, mammals and archelosaurians (turtles, crocodiles and birds; Fig 5b). Interestingly, a pineal or parietal eye is a primitive feature of bony fishes, since the parietal foramen, where the parietal eye sits, is an ancestral character (plesiomorphy) present in the fossilised crania of ancient fishes belonging to the ray- and lobe-finned lineages. Thus, parietal foramina are found in late Silurian bony fish like *Guiyu* and *Psarolepis* (Yu, 1998; Zhu et al, 1999; Zhu et al, 2009), early Devonian sarcopterygian fish like *Ligulalepis* and *Styloichthys* (Zhu and Yu, 2002; Clement et al, 2018), Devonian actinopterygians like *Meemannia, Mimipiscis* and *Raynerius* (Zhu et al. 2006; Choo 2011; Giles et al. 2015; Lu et al. 2016) as well as late Devonian tetrapodomorphs like *Ventastega, Tiktaalik, Acanthostega* and *Ichtyostega* (Daeschler et al. 2006; Ahlberg et al. 2008). Much later, during the Triassic and Jurassic, tetrapod lineages started to lose parietal foramina from the centre of the skull, as has been documented in detail for the therapsid lineage leading to the first mammals (Benoit et al. 2016). Thus, early sarcopterygians and tetrapods had both a parietal eye and *OPNLEP*, and it is tempting to speculate that, as the parietal eye was lost in later lineages, the *OPNLEP* gene faced the same fate (Fig 5b).

In conclusion, we have outlined the evolution of nonvisual OPN1 opsins in lepidosaurians and discovered a new opsin gene in this group. Our results imply a functional relation between the genomic repertoire of opsin genes and the parietal eye and habits of lizards, and as more genomic information accumulates and more species are represented, the details of nonvisual opsin evolution in this highly diverse clade will become clearer. As for OPNLEP, future studies are needed to address its photochemical characteristics, expression sites and physiological roles, in particular in relation to the reptilian third eye.

## Materials and Methods

### Genomic Database Searches

We used databases containing complete genome sequences of squamates and the tuatara, namely Ensembl (http://ensembl.org/index.html/), Ensembl Rapid Release (https://rapid.ensembl.org/index.html/) and, specially, the NCBI Genome Database (https://www.ncbi.nlm.nih.gov/datasets/genome/) (supplemental table S4). Initially, the well-characterised predicted protein sequences of OPNP, OPNPP, OPNPT and OPNVA from *X. laevis* and *X. tropicalis* (Bertolesi et al. 2020) were used to find the corresponding genes in *Anolis carolinensis* and a few other lizards using the TBLASTN searching tool. Often, the lizard genes had been automatically annotated with misleading names (for instance ‘parapinopsin-like’ for *OPNVA*). Then, the protein sequences from lizards were used to systematically search the genomes lepidosaurians for orthologue genes with TBLASTN. Alignments of the protein sequences with CLUSTAL OMEGA (Madeira et al. 2022) were used to confirm the identify of each opsin gene. We found that the genomic context (microsynteny) around the opsin genes was conserved, a feature that also helped identify the genes. In the process of collecting the protein sequences of the nonvisual OPN1 genes, we identified a new member of the group, *Lepidopsin* (*OPNLEP*), and repeated the analysis with this gene. The OPNLEP protein sequences analysed are compiled in Supplemental table S5.

### Assessing the Functionality of Opsin Genes

The integrity and functionality of the four previously known nonvisual opsin genes were evaluated by verifying that all exons could be retrieved from the genomes (four exons for *OPNPP* and *OPNPT*, five exons for *OPNP* and *OPNVA*) and that the whole protein sequence was predicted without frameshifts. For *OPNLEP*, which is a new gene without a standard sequence, we analysed in detail the *OPNLEP* loci from diverse lepidosaurians, namely *S. punctatus, A. carolinensis, P. vitticeps, P. muralis, V. komodensis* and *H. capensis*, to identify the precise boundaries of the four exons with their splicing sites and the coding sequence of the mRNAs.

To help in identifying exons, we used the global nucleotide alignment program MultiPipMaker (Schwartz et al. 2000; http://pipmaker.bx.psu.edu/pipmaker/). A 43 kb region encompassing the *OPNLEP* gene of *A. carolinensis*, including neighbouring genes *TNNC1* and *RPL29*, was used as default in MultiPipMaker to be compared to the homologous loci of other species and identify coding exons and splice sites. When the predicted protein sequences of opsins could not align smoothly with well-characterised proteins, as in the case of *OPNLEP* of *R. floridana* and *H. charlesbogerti*, we performed MultiPipmaker analyses with nucleotide sequences and visually inspected the alignments to identity indels, frameshifts and mutations in splice sites, so as to confirm their categorisation as pseudogenes. Similar analyses were performed with *OPNP, OPNPP* and *OPNPT*. The probable pseudogenes identified and the mutations that they carry are listed in Supplemental Table S3.

MultiPipMaker analyses also allowed for the identification of remnants of *OPNLEP* exons, as in geckos, snakes and other reptiles and vertebrates.

### Morphological and Activity Data

The presence of a pineal/parietal eye in lizards of particular species, genera and/or families was based on data collected by Gundy and Wurst (1976a, 1976b) and the character matrix built by Lee (1998), in which the presence of a pineal foramen is character number 33. For several species, a parietal foramen was observed in cranial scans collected by Watanabe et al (2019) and accessible on the Phenome10k website (http://phenome10k.org/), as well as the general scientific literature on anatomy. When data on a particular species was not found, we considered as valid the information on the same genus or family. In general, the presence of a parietal foramen was equaled to the presence of a parietal eye (Edinger 1955). The only exception that we considered were the fossorial, legless Californian Anniellidae lizards, which have a well-developed parietal eye without a corresponding foramen (Gundy and Wurst 1976a, 1976b).

Data on the daily activity patterns of lizards (diurnal, nocturnal, cathemeral) were drawn from the SquamBase database (https://datadryad.org/stash/dataset/doi:10.5061/dryad.76hdr7t3b) compiled by Meiri (2024).

### Phylogenetic Analysis

The evolutionary relationships among complete nonvisual OPN1 sequences were performed using the NGPhylogeny program package (Lemoine et al. 2019; https://https://ngphylogeny.fr/). Protein sequences were aligned with CLUSTAL OMEGA and ambiguously aligned regions and sites were removed with Gblocks (Talavera & Castresana, 2007). The final protein alignment has 258 aminoacid positions. Phylogenetic reconstruction was inferred using the balanced minimum evolution method as implemented in the FastME 2.0 program (Lefort et al. 2015). The model of aminoacid substitution chosen was LG (Le and Gascuel. 2008) with gamma distribution rates across sites (parameter set to 1.0). Tree refinement was done by the Subtree Pruning and Regrafting (SPR) method and bootstraping with 1000 replicates were used to evaluate statistical robustness. The final tree was visualised using iTOL (Letunic and Bork 2024; Interactive Tree of Life, https://itol.embl.de/). The five groups of nonvisual OPN1 opsins (OPNP, OPNPP, OPNPT, OPNLEP and OPNVA) were also retrieved in trees inferred with Maximum Likelihood and Bayesian methods implemented in the NGPhylogeny package (data not shown). The OPNLEP protein sequences analysed are compiled in Supplemental table S5.

## Acknowledgements

This work was supported by Agencia Nacional de Promoción Científica y Tecnológica (ANPCyT), Argentina [grant number PICT-2020-SERIEA-03451] and by Consejo Nacional de Investigaciones Científicas y Técnicas (CONICET), Argentina [grant number PIP 11220200100296]. RR is a PhD fellow of CONICET.

## References

Adler K. 1976. Extraocular photoreception in amphibians. Photochem Photophysiol. 23(4):275–98. 10.1111/j.1751-1097.1976.tb07250.x

Ahlberg PE, Clack JA, Luksevics E, Blom H, Zupiņs I. 2008. Ventastega curonica and the origin of tetrapod morphology. Nature. 453(7199):1199–204. doi:10.1038/nature06991

Amemiya CT et al. 2013. The African coelacanth genome provides insights into tetrapod evolution. Nature 496(7445):311–316. doi:10.1038/nature12027

Beaudry FEG, Iwanicki TW, Mariluz BRZ, Darnet S, Brinkmann H, Schneider P, Taylor JS. 2017. The non-visual opsins: eighteen in the ancestor of vertebrates, astonishing increase in ray-finned fish, and loss in amniotes. J Exp Zool B: Mol Dev Evol 328(7): 685–696. 10.1002/jez.b.22773

Beltrami G, Bertolucci C, Parretta A, Petrucci F, Foà A. 2010. A sky polarization compass in lizards: the central role of the parietal eye. J Exp Biol. 213(Pt 12):2048–54. doi:10.1242/jeb.040246

Benoit J, Abdala F, Manger PR, Rubidge BS. 2010. The sixth sense in mammalian forerunners: Variability of the parietal foramen and the evolution of the pineal eye in South African Permo-Triassic eutheriodont therapsids. Acta Palaeont Pol. 61(4):777–789 10.4202/app.00219.2015

Bertolesi GE, Atkinson-Leadbeater K, Mackey EM, Song YN, Heyne B, McFarlane S. 2020. The regulation of skin pigmentation in response to environmental light by pineal Type II opsins and skin melanophore melatonin receptors. J Photochem Photobiol B. 212:112024. doi:10.1016/j.jphotobiol.2020.112024

Blackshaw S, Snyder SH. 1997. Parapinopsin, a novel catfish opsin localized to the parapineal organ, defines a new gene family. J Neurosci. 17(21):8083–92. doi:10.1523/JNEUROSCI.17-21-08083.1997

Brazeau MD, Friedman M. 2015. The origin and early phylogenetic history of jawed vertebrates. Nature. 520(7548):490–7. doi:10.1038/nature14438

Brownstein CD, MacGuigan DJ, Kim D, Orr O, Yang L, David SR, Kreiser B, Near TJ. 2024. The genomic signatures of evolutionary stasis. Evolution. 78(5):821–834. doi:10.1093/evolut/qpae028

Burbrink FT, Grazziotin FG, Pyron RA, Cundall D, Donnellan S, Irish F, Keogh JS, Kraus F, Murphy RW, Noonan B, Raxworthy CJ, Ruane S, Lemmon AR, Lemmon EM, Zaher H. 2020. Interrogating genomic-scale data for Squamata (lizards, snakes, and amphisbaenians) shows no support for key traditional morphological Relationships. Syst Biol. 69(3):502–520. doi:10.1093/sysbio/syz062

Cavin L, Guinot G. 2014. Coelacanths as “almost living fossils”. Front Ecol Evol 2:49. doi:10.3389/fevo.2014.00049

Choo. 2011. Revision of the actinopterygian genus Mimipiscis (=Mimia) from the Upper Devonian Gogo Formation of Western Australia and the interrelationships of the early Actinopterygii. Earth Environ Sci Trans. 102:77–104. 10.1017/S1755691011011029

Clement AM, King B, Giles S, Choo B, Ahlberg PE, Young GC, Long JA. 2018. Neurocranial anatomy of an enigmatic Early Devonian fish sheds light on early osteichthyan evolution. Elife. 7:e34349. doi:10.7554/eLife.34349

Clement AM, Cloutier R, Lee MSY, King B, Vanhaesebroucke O, Bradshaw CJA, Dutel H, Trinajstic K, Long JA. 2024. A Late Devonian coelacanth reconfigures actinistian phylogeny, disparity, and evolutionary dynamics. Nat Commun. 15(1):7529. doi:10.1038/s41467-024-51238-4

Daeschler EB, Shubin NH, Jenkins FA Jr. 2006. A Devonian tetrapod-like fish and the evolution of the tetrapod body plan. Nature. 440(7085):757–63. doi:10.1038/nature04639

Dendy A. 1911. On the structure, development and morphological interpretation of the pineal organs and adjacent parts of the brain in the tuatara (Sphenodon punctatus). Philos Trans R Soc Lond B Biol Sci 201:226–331. 10.1098/rstb.1911.0006

Dodt E. 1973. The Parietal Eye (Pineal and Parietal Organs) of Lower Vertebrates. In: Jung R. (eds) Visual Centers in the Brain. Handbook of Sensory Physiology, vol 7/3/3B. Springer, Berlin, Heidelberg. 10.1007/978-3-642-65495-4_4

Eakin RM. 1973. The Third Eye. University of California Press. 10.1525/9780520326323

Edinger T. 1955. The size of parietal foramen and organ in reptiles. A rectification. Bull Mus Comp Zool. 114:1–34. https://www.biodiversitylibrary.org/item/26724

Ellis-Quinn BA, Simony CA (1991) Lizard homing behavior: the role of the parietal eye during displacement and radio-tracking, and time-compensated celestial orientation in the lizard Sceloporus jarrovi. Behav Ecol Sociobiol 28:397–407. 10.1007/BF00164121

Ekström P, Meissl H. 2003. Evolution of photosensory pineal organs in new light: the fate of neuroendocrine photoreceptors. Philos Trans R Soc Lond B Biol Sci. 358(1438):1679–700. doi:10.1098/rstb.2003.1303

Emerling CA. 2017a. Archelosaurian color vision, parietal eye loss, and the crocodylian nocturnal bottleneck. Mol Biol Evol 34(3): 666–676. 10.1093/molbev/msw265

Emerling CA. 2017b. Genomic regression of claw keratin, taste receptor and light-associated genes provides insights into biology and evolutionary origins of snakes. Mol Phylogenet Evol. 115:40–49. doi:10.1016/j.ympev.2017.07.014

Engbretson GA, Hutchison VH. 1976. Parietalectomy and thermal selection in the lizard Sceloporus magister. J Exp Zool. 198(1):29–38. doi:10.1002/jez.1401980105

Engbretson GA, Reiner A, Brecha N. 1981. Habenular asymmetry and the central connections of the parietal eye of the lizard. J Comp Neurol. 198(1):155–65. doi:10.1002/cne.901980113

Frigato E, Vallone D, Bertolucci C, Foulkes NS. 2006. Isolation and characterization of melanopsin and pinopsin expression within photoreceptive sites of reptiles. Naturwissenschaften. 93(8):379–85. doi:10.1007/s00114-006-0119-9

Gans C, Montero R. 2008. An atlas of amphisbaenian skull anatomy. In book: Biology of the Reptilia, vol. 21, Morphology I, The Skull and Appendicular Locomotor Apparatus of Lepidosauria. Publisher: Society for the Study of Amphibians and Reptiles. Editors: Gans C, Gaunt AS, Adler K. ISBN 978-0-91698-476-2

Giles S, Darras L, Clément G, Blieck A, Friedman M. 2015. An exceptionally preserved Late Devonian actinopterygian provides a new model for primitive cranial anatomy in ray-finned fishes. Proc Biol Sci. 282(1816):20151485. doi:10.1098/rspb.2015.1485

Goicoechea N, Frost DR, De la Riva I, Pellegrino KCM, Sites J Jr, Rodrigues MT, Padial JM. 2016. Molecular systematics of teioid lizards (Teioidea/Gymnophthalmoidea: Squamata) based on the analysis of 48 loci under tree-alignment and similarity-alignment. Cladistics. 32(6):624–671. doi:10.1111/cla.12150

Gundy GC, Wurst GZ. 1976a. Parietal eye-pineal morphology in lizards and its physiological implications. Anat Rec. 185(4):419–31. doi:10.1002/ar.1091850404

Gundy GC, Wurst GZ. 1976b. The Occurrence of Parietal Eyes in Recent Lacertilia (Reptilia). J Herpetol. 10(2):113–121. 10.2307/1562791

Hagen JFD, Roberts NS, Johnston RJ Jr. 2023. The evolutionary history and spectral tuning of vertebrate visual opsins. Dev Biol. 493:40–66. doi:10.1016/j.ydbio.2022.10.014

Hankins MW, Davies WIL, Foster RG. 2014. The Evolution of Non-visual Photopigments in the Central Nervous System of Vertebrates. In: Hunt D, Hankins M, Collin S, Marshall N. (eds) Evolution of Visual and Non-visual Pigments. Springer Series in Vision Research, vol 4. Springer, Boston, MA. 10.1007/978-1-4614-4355-1_3

Hsiang AY, et al. 2015. The origin of snakes: revealing the ecology, behavior, and evolutionary history of early snakes using genomics, phenomics, and the fossil record. BMC Evol Biol. 15:87. doi:10.1186/s12862-015-0358-5

Hutchison VH, Kosh RJ. 1974. Thermoregulatory function of the parietal eye in the lizard Anolis carolinensis. Oecologia. 16:173–177. 10.1007/BF00345581

Labra A, Voje KL, Seligmann H, Hansen TF. 2010. Evolution of the third eye: a phylogenetic comparative study of parietal-eye size as an ecophysiological adaptation in Liolaemus lizards. Biol J Linn Soc. 101:870–83. doi:10.1111/j.1095-8312.2010.01599.x

Lamb TD. 2013. Evolution of phototransduction, vertebrate photoreceptors and retina. Prog Retin Eye Res. 36:52–119. doi:10.1016/j.preteyeres.2013.06.001

Lagman D, Bergqvist CA, Kuraku S. 2024. How did vertebrate visual opsins diversify? -putting the last pieces of the puzzle together. bioRxiv 2024.02.06.579127; 10.1101/2024.02.06.579127

Le SQ, Gascuel O. 2008. An improved general amino acid replacement matrix. Mol Biol Evol. 25(7):1307–20. doi:10.1093/molbev/msn067

Lee MSY. 1998. Convergent evolution and character correlation in burrowing reptiles: towards a resolution of squamate relationships. Biol J Linn Soc. 65(4):369–453. 10.1111/j.1095-8312.1998.tb01148.x

Lefort V, Desper R, Gascuel O. 2015. FastME 2.0: A Comprehensive, Accurate, and Fast Distance-Based Phylogeny Inference Program. Mol Biol Evol. 32(10):2798–800. doi:10.1093/molbev/msv150

Lemoine F, Correia D, Lefort V, Doppelt-Azeroual O, Mareuil F, Cohen-Boulakia S, Gascuel O. 2019. NGPhylogeny.fr: new generation phylogenetic services for non-specialists. Nucleic Acids Res. 47(W1):W260–W265. doi:10.1093/nar/gkz303

Letunic I, Bork P. 2024. Interactive Tree of Life (iTOL) v6: recent updates to the phylogenetic tree display and annotation tool. Nucleic Acids Res. 52(W1):W78–W82. doi:10.1093/nar/gkae268

Lu J, Giles S, Friedman M, den Blaauwen JL, Zhu M. 2016. The Oldest Actinopterygian Highlights the Cryptic Early History of the Hyperdiverse Ray-Finned Fishes. Curr Biol. 26(12):1602–1608. doi:10.1016/j.cub.2016.04.045

Katti C, Stacey-Solis M, Coronel-Rojas NA, Davies WIL. 2019. The diversity and adaptive evolution of visual photopigments in reptiles. Front Ecol Evol. 7:352. 10.3389/fevo.2019.00352

Kawano-Yamashita E, Koyanagi M, Terakita A. 2014. The Evolution and Diversity of Pineal and Parapineal Photopigments. In: Hunt D, Hankins M, Collin S, Marshall N. (eds) Evolution of Visual and Non-visual Pigments. Springer Series in Vision Research, vol 4. Springer, Boston, MA. 10.1007/978-1-4614-4355-1_1

Kojima D, Mano H, Fukada Y. 2000. Vertebrate ancient-long opsin: a green-sensitive photoreceptive molecule present in zebrafish deep brain and retinal horizontal cells. J Neurosci. 20(8):2845–51. doi:10.1523/JNEUROSCI.20-08-02845.2000

Kojima K, Matsutani Y, Yanagawa M, Imamoto Y, Yamano Y, Wada A, Shichida Y, Yamashita T. 2021. Evolutionary adaptation of visual pigments in geckos for their photic environment. Sci Adv. 7(40):eabj1316. doi:10.1126/sciadv.abj1316

Korf HW, Liesner R, Meissl H, Kirk A. 1981. Pineal complex of the clawed toad, Xenopus laevis Daud.: structure and function. Cell Tissue Res. 216(1):113–30. doi:10.1007/BF00234548

Koyanagi M, Kawano E, Kinugawa Y, Oishi T, Shichida Y, Tamotsu S, Terakita A. 2004. Bistable UV pigment in the lamprey pineal. Proc Natl Acad Sci U S A. 101(17):6687–91. doi:10.1073/pnas.0400819101

Koyanagi M, Wada S, Kawano-Yamashita E, Hara Y, Kuraku S, Kosaka S, Kawakami K, Tamotsu S, Tsukamoto H, Shichida Y, Terakita A. 2015. Diversification of non-visual photopigment parapinopsin in spectral sensitivity for diverse pineal functions. BMC Biol. 13:73. doi:10.1186/s12915-015-0174-9

Madeira F, Madhusoodanan N, Lee J, Eusebi A, Niewielska A, Tivey ARN, Meacham S, Lopez R, Butcher S. 2024. Using EMBL-EBI Services via Web Interface and Programmatically via Web Services. Curr Protoc. 4(6):e1065. doi:10.1002/cpz1.1065

Mano H, Fukada Y. 2007. A median third eye: pineal gland retraces evolution of vertebrate photoreceptive organs. Photochem Photobiol. 83(1):11–8. doi:10.1562/2006-02-24-IR-813

Max M, McKinnon PJ, Seidenman KJ, Barrett RK, Applebury ML, Takahashi JS, Margolskee RF. 1995. Pineal opsin: a nonvisual opsin expressed in chick pineal. Science. 267(5203):1502–6. doi:10.1126/science.7878470

Meiri S. 2024. SquamBase—A database of squamate (Reptilia: Squamata) traits. Glob Ecol Biogeogr. 33:e13812. 10.1111/geb.13812

Meiniel A. 1977. Morphogenèse du complexe pariétal de Lacerta vivipara J. Considérations sur l’origine du 3ème œil des lacertiliens. J. Neural Transmission. 41:287–311. 10.1007/BF01252023

Okano T, Yoshizawa T, Fukada Y. 1994. Pinopsin is a chicken pineal photoreceptive molecule. Nature. 372(6501):94–7. doi:10.1038/372094a0

Oksche A. 1965. Survey of the development and comparative morphology of the pineal organ. Prog Brain Res. 10:3–29. doi:10.1016/s0079-6123(08)63445-7

Papenfuss TJ, Parham JF. 2013. Four New Species of California Legless Lizards (Anniella). Breviora. 536(1):1–17. 10.3099/MCZ10.1

Pérez JH, Tolla E, Dunn IC, Meddle SL, Stevenson TJ. 2019. A Comparative Perspective on Extra-retinal Photoreception. Trends Endocrinol Metab. 30(1):39–53. doi:10.1016/j.tem.2018.10.005

Philp AR, Garcia-Fernandez JM, Soni BG, Lucas RJ, Bellingham J, Foster RG. 2000. Vertebrate ancient (VA) opsin and extraretinal photoreception in the Atlantic salmon (Salmo salar). J Exp Biol. 203(Pt 12):1925–36. doi:10.1242/jeb.203.12.1925

Pinto BJ, Nielsen SV, Gamble T. 2019. Transcriptomic data support a nocturnal bottleneck in the ancestor of gecko lizards. Mol Phylogenet Evol 141:106639. 10.1016/j.ympev.2019.106639.

Pyron RA, Burbrink FT, Wiens JJ. 2013. A phylogeny and revised classification of Squamata, including 4161 species of lizards and snakes. BMC Evol Biol. 13:93. doi:10.1186/1471-2148-13-93

Redmond AK, Casey D, Gundappa MK, Macqueen DJ, McLysaght A. 2023. Independent rediploidization masks shared whole genome duplication in the sturgeon-paddlefish ancestor. Nat Commun. 14(1):2879. doi:10.1038/s41467-023-38714-z

Sakai K, Imamoto Y, Su CY, Tsukamoto H, Yamashita T, Terakita A, Yau KW, Shichida Y. 2012. Photochemical nature of parietopsin. Biochemistry. 51(9):1933–41. doi:10.1021/bi2018283

Sato K, Yamashita T, Kojima K, Sakai K, Matsutani Y, Yanagawa M, Yamano Y, Wada A, Iwabe N, Ohuchi H, et al. 2018. Pinopsin evolved as the ancestral dim-light visual opsin in vertebrates. Commun Biol 1(1):1–10. 10.1038/s42003-018-0164-x

Schwartz S, Zhang Z, Frazer KA, Smit A, Riemer C, Bouck J, Gibbs R, Hardison R, Miller W. 2000. PipMaker--a web server for aligning two genomic DNA sequences. Genome Res. 10(4):577–86. doi:10.1101/gr.10.4.577

Simões BF, Sampaio FL, Jared C, Antoniazzi MM, Loew ER, Bowmaker JK, Rodriguez A, Hart NS, Hunt DM, Partridge JC, Gower DJ. 2015. Visual system evolution and the nature of the ancestral snake. J Evol Biol. 28(7):1309–20. doi:10.1111/jeb.12663

Simões TR, Vernygora O, Caldwell MW, Pierce SE. 2020. Megaevolutionary dynamics and the timing of evolutionary innovation in reptiles. Nat Commun. 11(1):3322. doi:10.1038/s41467-020-17190-9

Simões TR, Pyron RA. 2021. The Squamate Tree of Life. Bull Mus Comp Zool. 163(2):47–95. 10.3099/0027-4100-163.2.47

Simões TR, Kammerer CF, Caldwell MW, Pierce SE. 2022. Successive climate crises in the deep past drove the early evolution and radiation of reptiles. Sci Adv. 8(33):eabq1898. doi:10.1126/sciadv.abq1898.

Smith KT, Bhullar BS, Köhler G, Habersetzer J. 2018. The Only Known Jawed Vertebrate with Four Eyes and the Bauplan of the Pineal Complex. Curr Biol. 28(7):1101-1107.e2. doi:10.1016/j.cub.2018.02.021

Solessio E, Engbretson GA. 1993. Antagonistic chromatic mechanisms in photoreceptors of the parietal eye of lizards. Nature. 364(6436):442–5. doi:10.1038/364442a0

Stebbins RC, Eakin RM. 1958. The role of the “third eye” in reptilian behavior. Am Mus Novit. 1870:1–40. http://hdl.handle.net/2246/4659

Su CY, Luo DG, Terakita A, Shichida Y, Liao HW, Kazmi MA, Sakmar TP, Yau KW. 2006. Parietal-eye phototransduction components and their potential evolutionary implications. Science. 311(5767):1617–21. doi:10.1126/science.1123802

Talavera G, Castresana J. 2007. Improvement of phylogenies after removing divergent and ambiguously aligned blocks from protein sequence alignments. Syst Biol. 56(4):564–77. doi:10.1080/10635150701472164

Taniguchi Y, Hisatomi O, Yoshida M, Tokunaga F. 2001. Pinopsin expressed in the retinal photoreceptors of a diurnal gecko. FEBS Lett 496(2–3):69–74. 10.1016/S0014-5793(01)02395-X

Terakita A. 2005. The opsins. Genome Biol. 6(3):213. 10.1186/gb-2005-6-3-213

Terakita A, Kawano-Yamashita E, Koyanagi M. 2012. Evolution and diversity of opsins. WIREs Membr Transp Signal. 1:104–111. doi:10.1002/wmts.6

Traeholt C. 1997. Effect of masking the parietal eye on the diurnal activity and body temperature of two sympatric species of monitor lizards, Varanus s. salvator and Varanus b. nebulosus. J Comp Physiol B 167:177–184. 10.1007/s003600050062

Trost E. 1956. Über die Lage des Foramen Parietale bei rezenten Reptilien und Labyrinthodontia. Acta Anat. 26(4):318–339. 10.1159/000141106

Toriño P, Soto M, Perea D. 2021. A comprehensive phylogenetic analysis of coelacanth fishes (Sarcopterygii, Actinistia) with comments on the composition of the Mawsoniidae and Latimeriidae: evaluating old and new methodological challenges and constraints. Hist Biol. 33(12):3423–3443. 10.1080/08912963.2020.1867982

Tosini G. 1997. The pineal complex of reptiles: physiological and behavioral roles. Ethol Ecol Evol. 9(4):313–333. 10.1080/08927014.1997.9522875

Ung CY, Molteno AC. 2004. An enigmatic eye: the histology of the tuatara pineal complex. Clin Exp Ophthalmol. 32(6):614–8. doi:10.1111/j.1442-9071.2004.00912.x

van de Kamer JC, Feekes C, Burgers AC. 1962. Histological investigation of the unpigmented meningeal spot on the brain of black background adapted Xenopus laevis larvae. Z Zellforsch Mikrosk Anat. 56:359–70. doi:10.1007/BF00334793

Vigh B, Manzano MJ, Zádori A, Frank CL, Lukáts A, Röhlich P, Szél A, Dávid C. 2002. Nonvisual photoreceptors of the deep brain, pineal organs and retina. Histol Histopathol. 17(2):555–90. doi:10.14670/HH-17.555

Wada S, Kawano-Yamashita E, Koyanagi M, Terakita A. 2012. Expression of UV-sensitive parapinopsin in the iguana parietal eyes and its implication in UV-sensitivity in vertebrate pineal-related organs. PLoS One. 7(6):e39003. doi:10.1371/journal.pone.0039003

Watanabe A, Fabre AC, Felice RN, Maisano JA, Müller J, Herrel A, Goswami A. 2019. Ecomorphological diversification in squamates from conserved pattern of cranial integration. Proc Natl Acad Sci U S A. 116(29):14688–14697. doi:10.1073/pnas.1820967116

Yu X. 1998. A new porolepiform-like fish, Psarolepis romeri, gen. et sp. nov. (Sarcopterygii, Osteichthyes) from the Lower Devonian of Yunnan, China. J Verteb Paleontol. 18(2):261–274, DOI:10.1080/02724634.1998.10011055

Zhu M, Yu X, Janvier P. 1999. A primitive fossil fish sheds light on the origin of bony fishes. Nature. 397:607– 610. 10.1038/17594

Zhu M, Yu X. 2002. A primitive fish close to the common ancestor of tetrapods and lungfish. Nature. 418(6899):767–70. doi:10.1038/nature00871

Zhu M, Yu X, Wang W, Zhao W, Jia L. 2006. A primitive fish provides key characters bearing on deep osteichthyan phylogeny. Nature. 441(7089):77–80. doi:10.1038/nature04563

Zhu M, Zhao W, Jia L, Lu J, Qiao T, Qu Q. 2009. The oldest articulated osteichthyan reveals mosaic gnathostome characters. Nature. 458(7237):469–74. doi:10.1038/nature07855

