## Supplemental Figures S1 & S2 for "Evolution of pineal non-visual opsins in lizards and the tuatara (Lepidosauria)"

```
OPNLEP  ------------MMNHS-------LFSSNALTFLGPNATCNSTTCLDQASEKKYKF-GVG
OPNPT   ------------MENESS-LVLEGAEGYIVRPTIFPR----------------AGYGVLA
OPNPP   -----MDSLNA---------STLSSNASTVRVVLMPR----------------IGYTIIA
OPNP    -----------MWVSNEWINGTPGPFDGPQWPFLAPRGT-----------Y-----TSVA
VA      MSKPDMDGLRREAENDSWLFN-PSSSSAPFDPFLQPLDI-----------IEPWNFHLIS
                                          : *                     :.
                TM1                                    TM2
OPNLEP  VLMILGLFLGFWFNCIIVILTIRHKFLRNPINYAMVNLSVADMFSIVFGLGPTMDANLKG
OPNPT   FLMFINALFSLFNNFLVIAVTLKNPQLRNPINIFILNLSFSDLMMSICGTTIVIATNYHG
OPNPP   IIMATSCTLSVILNTAVIAVTIKYRQLRQPINYSLVNLAIADLGAALLGGSLNVETNAVG
OPNP    VLMGLVMVSAAVVNGLVILVSVQYKRLRSPLNYILVNLAVADLLVTFFGSTISFANNIYG
VA      ALMFVVTLLSLSENFTVILVTIKFKQLRQPLNYVIVNLSVADFLVSLIGGTISFSTNLKG
          :*       .   *  :: .:.:    **.*:*  ::**:.:*:    .  *     *  *
                           TM3                                  TM4
OPNLEP  YSFQSKSFCAFQGFCITLFGLVALWSVVLCAVERYLLVCRPFKALHFGKRHAVLALLFIW
OPNPT   YFYLGRRFCIFQGFAVNYFGIVSLWSLTILAYERYNVVCQPLGTLQMSTQRAYQLLGFIW
OPNPP   YYNLGRVGCVTEGFAMAFFGIVALCTIAVIAVDRAIVIAKPMGTITFTTRKAMIGVAVSW
OPNP    FFVFGPTACEFEGFMVSLTGIVGLWSLAILAFERYLVICKPVGDFRFQQRHAVIGCAFTW
VA      YFYMGHWACVLEGFAVTFFGIVALWSLALLAFERYVVICRPLGNMRLKGKHAALGIAFVW
        :   .    *   :** :     *:*.* ::.: *  :*   ::.:*.   :  :  *  .  *
                                                        TM5
OPNLEP  LWSLIWTLPPLFGWSSYEVEGIASNCALAWHRKDWKSRSYHLSLFFVCFLVPFILMSLSY
OPNPT   VFCLFWAVVPLFGWSSYGPEGVQTSCSIGWEERSWNNYSYLIVYFLSCFFIPVLIIGFSY
OPNPP   IWSLVWNTPPLFGWGGYEMEGVMTSCAPDWANSDPINVSYIICYFLFCFTIPFITILVSY
OPNP    LWSLLWTLPPLFGWSSYIPEGLRTSCGPNWYTGGSDNNSYIMTLFVTCFVTPLAMIVFSY
VA      IFSFIWTVPPIMGWSSYTTSKIGTTCEPNWYSGDYNDHTFIITFFTTCFILPLLVILVSY
        ::.:.:.*   *::**..*   .  :  :.*    *    .:: :   *   **   *.   :  .**
                                         TM6
OPNLEP  GMLIFTLRKVAKTGLVQSNSTQRAESQ--VTKTAAIMILAFLLSWMPYAAFALTKVVKPD
OPNPT   GNVIRSLHGLNKK-VEQLGGKSNPEEEFRAVIMVLVMVVAFLICWLPYTLFALTVVFNPA
OPNPP   GYLIWTLRQVAKVGLAQXGSTTKAEAQ--VSRMVIVMVMAFLICWLPYATFALVVVGNPK
OPNP    ANLLLTLRAVAAQ-QKELETTQRAERE--VTRMVITMVMAFLVCWLPYASFAMVVATNKD
VA      GKLMRKLRKVSDT-QGRLGTTRKPERQ--VTRMVVIMILAFLICWSPYAAFSILVTTCPS
         .  ::  .*:  :           .  .  *  :    .     .    *::***:.*  **:  *::    .
                          TM7
OPNLEP  VTMDPNIQSIPLFMAKTGTIYNPVVYVCLNKQYREQVAASLRCLRKCRVVPSKHRVLNAL
OPNPT   LNISPLAATIPTYLSKTSPVYNPIIYIFLNKEFRECAVEFITCGKVVLTSPEEDISTSAI
OPNPP   IYINPIIATVPMYMAKSSTFYNPIIYIFMNKQFRDCLVRCLLCGRNPCASEQTDEDDLEV
OPNP    LFIQPALASLPSYFSKTATVYNPIIYVFMNKQFRSCLLSTMSCGRRPVAAQGTTPAMISS
VA      IELDPRLAAIPAFFSKTATVYNPVIYVFMNNQFRKCLVQLFRCSDQEIVDANVNPISEKD
         :  :.*     ::*  :::*:.  .***::*:  :*:::*.       :  *        .

OPNLEP  PPALKRLQGK--QLSCNTNRIAPQNE-----------LNISRNLLLPNMSYSDDESSSQ
OPNPT   SDEGIAPCK--------INQVTPV-------------------------------------
OPNPP   S----------------T--IAPAPPTRRGKVAPI-------------------------
OPNP    PRGLTS------ALEGSRNKVVPSASEGSGND--VGTS----------------------
VA      TLAHTKHCGEMSTVAAHVIVFNPRSEDQHGSCQTFAQLAISENKVYPV-----------
                            . *
```

**Supplemental Figure S1:** Clustal Omega alignment between predicted aminoacid sequences of nonvisual OPN1 opsins from *Anolis carolinensis* (green anole). Predicted transmembrane domains (TM) are highlighted.

Romero & de Souza, 2024

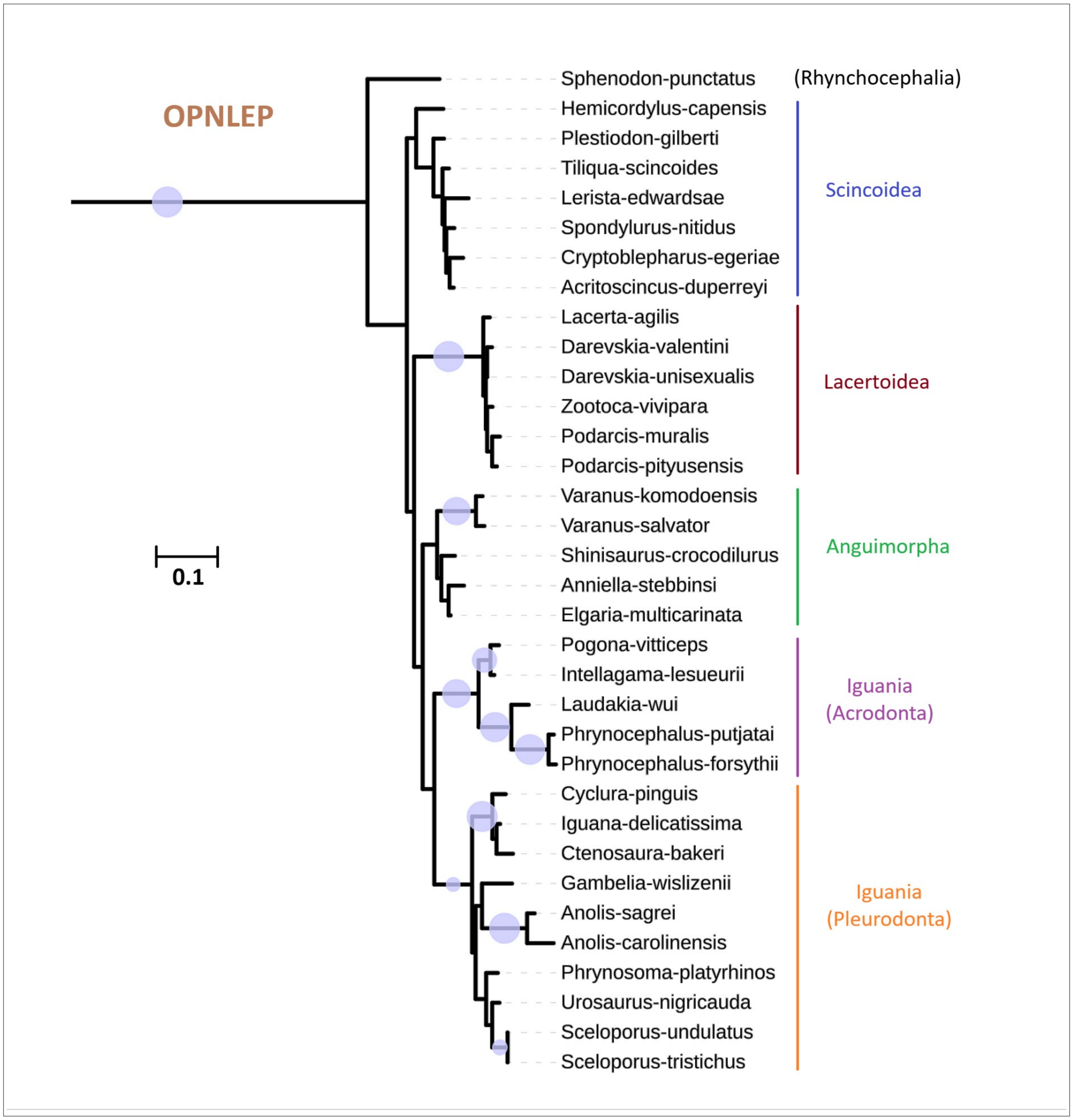


**Supplemental Figure S2:** Close-up of the OPNLEP branch of the phylogenetic tree shown in Figure 3 inferred with FastME 2.0 (balanced minimum evolution). OPNLEP proteins are grouped around known lepidosaurian clades. Blue circles indicate branches with >900 bootstrap values.

Romero & de Souza, 2024
